# Warming disrupts plant-fungal endophyte symbiosis more strongly in leaves than roots

**DOI:** 10.1101/2025.01.22.634355

**Authors:** Joseph D. Edwards, Melanie R. Kazenel, Yiqi Luo, Joshua S. Lynn, Rebecca L. McCulley, Lara Souza, Carolyn Young, Jennifer A. Rudgers, Stephanie N. Kivlin

## Abstract

Disruptions to species interactions from global change will negatively impact plant primary production, with broader consequences for species’ abundances, distribution, and community composition. Fungal endophytes that live inside plant leaves and roots could potentially mitigate plant heat stress from global warming. Conversely, disruptions of these symbioses could exacerbate the negative impacts of warming. To better understand the consistency and strength of warming-induced changes to fungal endophytes, we examined fungal leaf and root endophytes in three grassland warming experiments in the US ranging from 2 to 25 years and spanning 2000 km, 12 °C of mean annual temperature, and 600 mm of precipitation. We found that experimental warming disrupted symbiosis between plants and fungal endophytes. Colonization of plant tissues by septate fungi decreased in response to warming by 90% in plant leaves and 35% in roots. Warming also reduced fungal diversity and changed community composition in plant leaves but not roots. The strength, but not direction, of warming effects on fungal endophytes varied by up to 75% among warming experiments. Finally, warming decoupled fungal endophytes from host metabolism. Overall, warming-driven disruption of fungal endophyte community structure and function suggests that this symbiosis may not be a reliable mechanism to promote plant resilience and ameliorate stress responses under global change.

## Introduction

Predicting how organisms will respond and acclimate to global change, particularly to climate warming, is among the greatest challenge facing ecosystems over the next century (Love et al., 2023). In long-lived plants these acclimations could be via physiological responses of plants themselves (Reich et al., 2016) or through their relationships with other species, including microbial symbionts, such as fungal endophytes in leaves and roots (Kivlin and Rudgers, 2019, Rodriguez et al., 2008). Alternatively, warming could disrupt symbiotic interactions (Zhu et al., 2022), with potential declines in plant resilience to climate change. A framework of how and when plant-fungal symbiosis will respond to climate warming, and in turn affect plant performance, across plant hosts and environments is critical because many long-lived plant species will need to acclimatize to climate change in order to survive over the next century (Smith et al., 2009).

Fungal endophytes occur in every plant species surveyed to date (Harrison and Griffin, 2020). These symbionts vary in abundance and composition across environmental climate gradients (Davison et al., 2015, Giauque and Hawkes, 2013, Kivlin et al., 2011, Kivlin et al., 2017, Zimmerman and Vitousek, 2012), within climate manipulations (reviewed by Kivlin et al., 2013, Kazenel et al., 2019, Lyons et al., 2021), and over time with climate change (Giauque and Hawkes, 2016). For example, dark septate endophyte (DSE) abundance typically increases with environmental stress and is associated with increased plant tolerance to high temperatures (Reininger and Sieber, 2012, Slaughter et al., 2018), among other stressors (Kivlin et al., 2013, Rodriguez et al., 2009). The large environmental and plant-host-driven variation in fungal endophyte symbiont communities complicates creating a predictive framework for the functional outcome of plant-fungal symbiosis under global change (Johnson et al., 1997, Johnson et al., 2010). Despite the strong influence of environmental factors on fungal endophyte composition and function, variation in endophyte composition can be just as large among plant hosts (Arnold and Lutzoni, 2007), with differences due to plant evolutionary lineage (Öpik et al., 2010), physiology (Hetrick et al., 1990, Higgins et al., 2007), size (Kivlin et al., 2019), or other leaf or root morphological traits (Kembel and Mueller, 2014, Valkama et al., 2005). These multifaceted drivers of fungal endophytes, plants, and their symbioses need further investigation to understand the generalizability in response to warming across time and space.

Defining plant tissue-specific rules for fungal endophyte responses to environmental change will be crucial for predicting their future dynamics and functions. Most studies examine above- and below-ground fungal groups independently. However, when examined together under climate change contexts, phylosphere fungal abundance, diversity, and composition have typically been reported to be more sensitive to warming than fungi within roots (Coince et al., 2014, Kazenel et al., 2019). Similarly, across seasons, leaf endophytes were more sensitive to winter temperature extremes than root endophytes (Wearn et al., 2012). Soil buffering of atmospheric temperature extremes may minimize diurnal and annual temperature shifts experienced by fungi that reside belowground (Fujimura et al., 2008). To characterize the whole-plant microbiome response to global change, we must consider differences among plant tissues.

Host organisms and their symbionts may have coordinated or discordant responses to warming. Within an individual plant, warming can trigger many physiological and metabolic changes associated with heat stress regulation (Suzuki et al., 2012), water conservation (Berry and Bjorkman, 1980, Reich et al., 2016), growth phenology (Miller-Rushing and Primack, 2008), and resource allocation (Rustad et al., 2001). At the same time, diverse fungal species may each respond individually to warming, with changes to fungal physiology that cascade to reorder the relative abundances of taxa residing within plant tissues (Rudgers et al., 2020). Fungal endophytes can promote local adaptation in extreme environments by helping to regulate oxidative stress and osmotic control (Acuña-Rodríguez et al., 2024). Fungal endophytes may also represent emergent vectors for development of plant thermotolerance via epigenetic mechanisms (Gilbert et al., 2010, Carrell et al., 2022). Unfortunately, warming can also increase fungal endophyte community heterogeneity among plant species (Jiang et al., 2021) and disrupt molecular signaling between plants and their fungal symbionts (Binet et al., 2017). Proximal mechanisms behind the benefits or costs of symbionts to plants are difficult to parse in the complex phytobiome (Giauque et al., 2019) though new evidence points to fungal-produced or fungal-induced plant metabolites as a causal driver of plant stress acclimation (Connor et al., 2017, Zeilinger et al., 2016). Yet, we still do not know the degree to which fungal associations, or the metabolites they induce, are specialized across geographic space, time since climate warming, or different plant species, precluding generalizable mechanisms of plant thermal response in future climates. An improved understanding of the direction and fidelity of endophytic fungal symbiosis responses to warming is necessary to predict future patterns under global change.

To better understand the consistency and strength of warming-induced changes to fungal endophytes, we examined fungal leaf and root endophytes for eight plant species in three US air temperature warming experiments spanning 2000 km, 12 °C of mean annual temperature, and 600 mm of rainfall (Figure 1). Warming experiments were sampled at 2, 17, or 25 years after initiation and included perennial grass species that spanned both C_3_ and C_4_ physiologies, allowing us to assess generalizable patterns of fungal endophyte response to warming climate conditions. We addressed three questions: (1) Does warming disrupt relationships between plants and fungal endophytes by reducing colonization and diversity or altering community composition of fungal endophytes, and are effects greater for leaves than roots? We predicted that root fungal endophytes would be more buffered from heat than leaf fungal endophytes because of both the insulating properties of soil relative to air and the greater diversity of fungal taxa colonizing roots relative to leaves. (2) Does warming alter leaf and root fungal endophytes in similar ways across diverse environmental conditions? By leveraging three existing warming experiments, which varied in the duration of warming, plant community composition, geography, and climate, we aimed to determine the generalizability of warming impacts on fungal endophytes both above- and below-ground. (3) Do warming-induced changes to fungal endophyte communities correspond to altered host function via plant metabolic activity? We predicted that responses of plants and fungi to heat, such as plant stress metabolism and reordering in fungal symbiont relative abundances, would reduce the coordination between endophyte communities and plant metabolism that occurs under ambient climate conditions.

**Figure 1.**
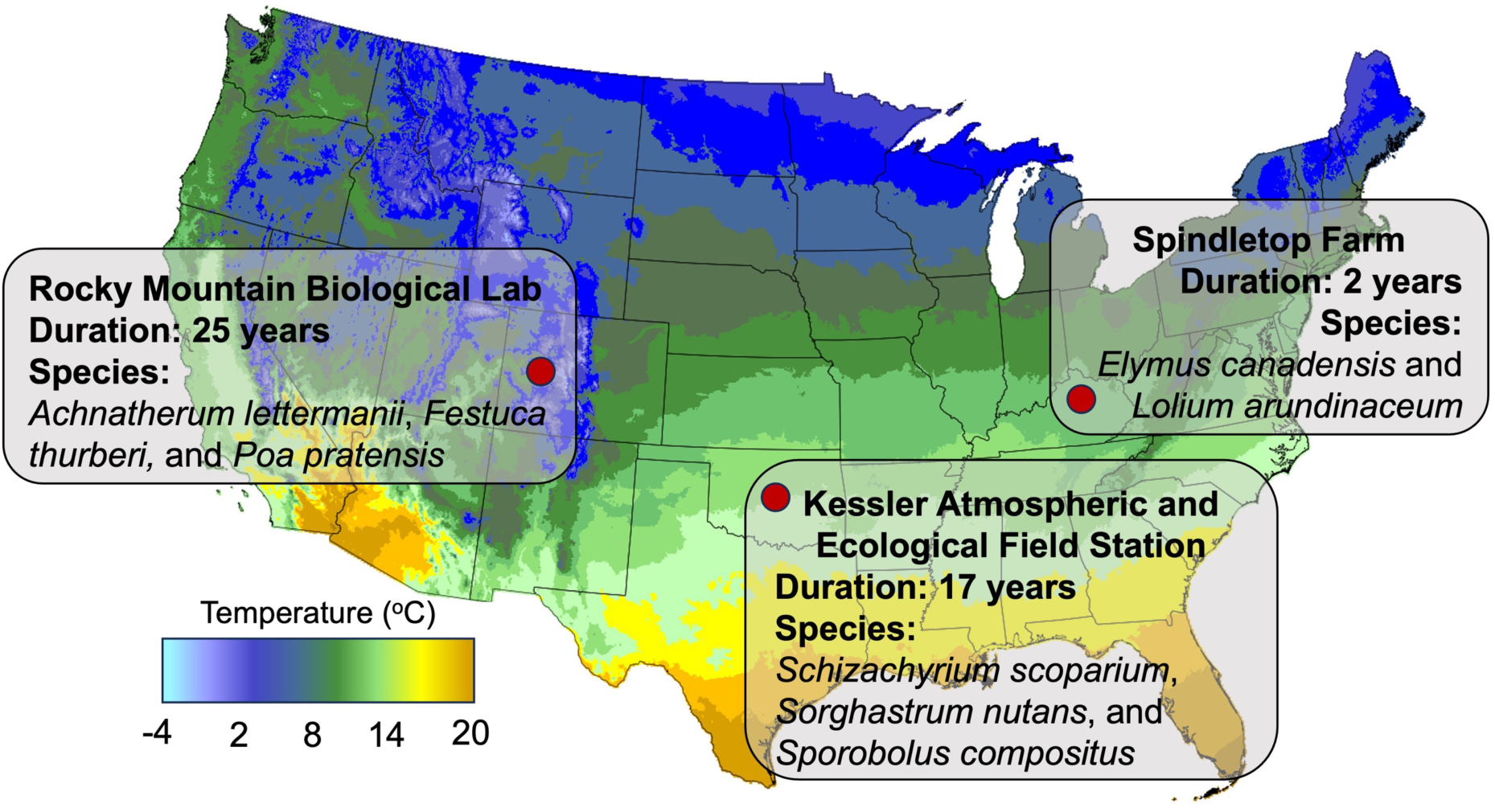
Location, age, and plant species from each warming experiment that experimentally warmed air plots with aboveground heaters, relative to ambient temperature controls. The Rocky Mountain Biological Laboratory (RMBL) experiment is conducted in Gothic, CO (38.95°N, 106.98°W) with an annual ambient temperature of 4.0 °C warmed by 2.1 °C. The Kessler Atmospheric and Ecological Field Station (Kessler) experiment was conducted in Washington, OK (35.06°N, 97.48°W) with an annual ambient temperature of 16 °C warmed by 1.2 °C. The Spindletop Farm (Spindletop) experiment was conducted in Lexington, KY (31.11°N, 84.49°W) with an ambient annual temperature of 12 °C, warmed by 3.0 °C.

## Methods

We sampled three grassland warming experiments across the United States, at the Rocky Mountain Biological Laboratory (hereafter RMBL) Colorado, Kessler Atmospheric and Ecological Field Station (hereafter Kessler) Oklahoma, and Spindletop Farm (hereafter Spindletop) Kentucky. Sites varied in ambient climate, species composition, location, and warming duration (Figure 1).

### Warming experimental designs

#### RMBL

We sampled one of the oldest active warming experiments in the world (at sampling), which was installed in 1991 at the RMBL. Electric heaters (22 W m^2^ infrared) were suspended 1.5 m above plots (10m x 3m) to simulate the soil surface heating expected under a doubling of CO_2_ (n = 5; Harte et al., 1995). Soil moisture and temperature were logged every 2 h. Heating warmed the top 15 cm of soil by ∼2 °C, dried it by 10-20% and prolonged the snow-free season at each end by ∼12 d.

#### Kessler

The Kessler warming experiment also used infrared heaters to warm plots (2 m x 2 m; 100 W m^2^, 1.5 m aboveground) since 1999 with soil temperature and moisture recorded hourly (n = 5). Warming treatments increased aboveground temperatures by 1.21 °C and soil temperature by 1.71°C in the top 10 cm, and decreased soil moisture by 1.60 % (Wan et al., 2002).

#### Spindletop

The Spindletop warming experiment heated plots (3 m diameter) with aboveground infrared heaters from 2009-2013, was harvested and then run again continuously since 2015, with soil moisture measured every 15 m (n = 5). Warming increased aboveground air temperature by 3.04 °C and decreased soil moisture by 13.9 % (McCulley et al., 2014).

#### Shared methods

All warming experiments were sampled during the growing season in 2016 (RMBL August, Kessler July, and Spindletop October). In each experiment, leaves and roots were collected separately for 3-6 individuals per plot from each of the dominant grass species and combined by plant tissue type for each species. At RMBL, plant species included *Achnatherum lettermanii*, *Festuca thurberi,* and *Poa pratensis*. Kessler plants sampled were *Schizachyrium scoparium*, *Sorghastrum nutans*, and *Sporobolus compositus*. *Elymus canadensis* and *Lolium arundinaceum* were collected from Spindletop. Spindletop plants were part of an experimental manipulation with and without leaf endophytes (*Epichloe spp.*). Thus, we considered endophyte-symbiotic (e+) and non-symbiotic (e-) plants as different species in our analysis. All grasses were perennial species; at RMBL and Spindletop grasses were C_3_ while the Kessler species were C_4_ in terms of photosynthetic pathway.

### Fungal symbiont colonization

To assess fungal colonization of grass roots and leaves, we stained tissue samples and scored colonization via light microscopy (Bacon and White, 2018, Mcgonigle et al., 1990). For leaf tissue, we calculated the hyphal length of all fungi in μm per mm^2^ leaf tissue. For root tissue, we separately quantified septate hyphal % colonization (*i.e.,* plant saprotrophs and pathogens; Ascomycota & Basidiomycota), aseptate hyphal % colonization (arbuscular mycorrhizal [AM] fungi; Glomeromycotina), and relative abundance of AM fungal functional structures including arbuscules and vesicles as % colonization.

### Fungal community composition

We surface-sterilized all plant material and stored it frozen at -20 °C until extraction. We then extracted fungal DNA from 0.25 g of frozen plant tissue with the Mo-Bio DNeasy plant mini kit (Qiagen, Germantown, MD, USA). DNA was quantified fluorometrically (Qubit, Invitrogen, Carlsbad, CA, USA) and normalized to 20 ng/µl for subsequent PCR. For PCR, we used Illumina TruSeq V3 indices (Illumina, San Diego, CA, USA) linked to ITS2 rDNA fungal-specific primers (5.8S-Fun/ITS4-Fun; Taylor et al., 2016). These primers allow for the detection of the largest swath of fungi while also restricting non-target taxa (e.g., plants and animals). Reactions contained 20.5 µl of platinum PCR Supermix (Invitrogen, Carlsbad, CA, USA), 1.25 µl of each primer (10 µM), 0.5 µl of BSA (20mg/mL), and 2 µl of DNA. All PCRs were performed in triplicate with a hot start at 94 °C for 3 min, and 25 cycles of 94 °C for 45s, 50 °C for 1 min, 72 °C for 90 s, and a final extension step of 72 °C for 10 min. Triplicate PCRs were combined and cleaned with Agencourt AMPure XP magnetic beads (Beckman Coulter, Brea, CA) and quantitated with a Qubit fluorometer. Samples were pooled in equal amounts and sequenced on Illumina MiSeq v3 (2 x 250b PE run) at the University of Texas Genome Sequencing and Analysis Facility (GSAF).

### Bioinformatics: Sequences

All fungal sequences were processed in QIIME v. 1.9.1 (Caporaso et al., 2010) using standard scripts to join paired end reads, remove any unjoined sequences, remove forward and reverse primers, and filter sequences with quality scores < 25. Chimeras were removed using UCHIME with default parameters (Edgar, 2010). We then used UCLUST (Edgar et al., 2011) to create operational taxonomic units (OTUs) at 97% identity and removed singleton OTUs. We chose to use 97% OTUs, as opposed to exact sequencing variants, as these likely provide more accurate estimates of diversity metrics due to conflation of inter- and intra-specific diversity without clustering (Kauserud, 2023, Lofgren et al., 2019). Moreover, ecological patterns of fungal assemblages are robust to classification techniques across ecosystems (Glassman and Martiny, 2018). To assign taxonomy we used a naive Bayesian classifier (Wang et al., 2007) with the UNITE fungal training set (v10.0) for ITS2 data (Abarenkov et al., 2023). Taxonomies assigned to at least the genus level were then annotated with functional assignations based on the FungalTraits database (Pölme et al., 2021), which we aggregated into mutualistic, pathogenic, and saprotrophic guilds. Sequences were deposited in the NCBI Sequence Read Archive (SRA) under (Bioproject PRJNA1138416 to be released upon manuscript acceptance). Samples containing fewer than 2500 fungal reads (*n* = 5) were discarded and all remaining OTU abundances were Hellinger transformed to equalize sampling effort, as proportional transformation is one of the most accurate ways to account for differential sampling effort in large microbial datasets (Labouyrie et al., 2023, McKnight et al., 2019).

### Metabolomics extraction methods and quantification

We performed metabolomic analyses for the RMBL and Kessler warming experiments but were unable to collect sufficient plant biomass from the Spindletop warming experiment for these analyses. Plant leaf and root samples (100 mg) were weighed into 1.5 mL centrifuge tubes. We added 1.3 mL of extraction solvent (40:40:20 HPLC grade methanol, acetonitrile, water with 0.1% formic acid) pre-chilled to 4 °C to each tube. Samples were vortexed to suspend non-plant particles and the extraction was allowed to proceed for 20 min at 4 °C while being shaken in an orbital platform shaker (Bellco, Vineland, NJ). The samples were centrifuged for 5 min (16.1 rcf) at 4 °C. The supernatant was transferred to new 1.5 mL centrifuge tubes and the plant tissue was resuspended with 50 µL of extraction solvent. Extraction was allowed to proceed for 20 min at 4 °C while being shaken. This step was repeated once more. The centrifuge tubes containing all collected supernatant liquid were centrifuged for 5 min (16.1 rcf) at 4 °C to remove any remaining soil particles and 1.2 mL were transferred to vials. Vials containing 1.2 mL of the collected supernatant were dried under a stream of N_2_ until all the extraction solvent had been evaporated. Solid residue was resuspended in 300 µL of sterile water and transferred to 300 µL autosampler vials. Samples were immediately placed in autosampler trays for mass spectrometry.

Samples placed in an autosampler tray were kept at 4 °C. A 10 µL aliquot was injected through a Synergi 2.5 µm reverse-phase Hydro-RP 100, 100 x 2.00 mm LC column (Phenomenex, Torrance, CA) kept at 25 °C. The eluent was introduced into the MS via an electrospray ionization source conjoined to an Exactive™ Plus Orbitrap Mass Spectrometer (Thermo Scientific, Waltham, MA) through a 0.1 mm internal diameter fused silica capillary tube. The mass spectrometer was run in full scan mode with negative ionization mode with a window from 85 – 1000 m/z. with a method adapted from Lu et al. (2010). The samples were run with a spray voltage was 3 kV. The nitrogen sheath gas was set to a flow rate of 10 psi with a capillary temperature of 320°C. AGC (acquisition gain control) target was set to 3e6. The samples were analyzed with a resolution of 140,000 and a scan window of 85 to 800 m/z for from 0 to 9 minutes and 110 to 1000 m/z from 9 to 25 minutes. Solvent A consisted of 97:3 water:methanol, 10 mM tributylamine, and 15 mM acetic acid. Solvent B was methanol. The gradient from 0 to 5 minutes is 0% B, from 5 to 13 minutes is 20% B, from 13 to 15.5 minutes is 55% B, from 15.5 to 19 minutes is 95% B, and from 19 to 25 minutes is 0% B with a flow rate of 200 µL/min.

### Bioinformatics: Metabolomics

Files generated by Xcalibur (RAW) were converted to the open-source mzML format (Martens et al. 2011) via the open-source msconvert software as part of the ProteoWizard package (Chambers et al., 2012). Maven (mzroll) software, Princeton University (Clasquin et al., 2012, Melamud et al., 2010) was used to automatically correct the total ion chromatograms based on the retention times for each sample. Metabolites were manually identified and integrated using known masses (± 5 ppm mass tolerance) and retention times (≤ 1.5 m), only identified metabolites were used for this analysis. Metabolomics data were transformed based on recommended practices (Sun and Xia, 2024) by adding 20% of the minimum non-zero value of each metabolite to account for minimum instrument read accuracy and unit-scaled to account for differences in the relative abundances of each metabolite. All subsequent analyses were performed in R (Team, 2013) unless noted otherwise.

### Data analysis: Abundance and diversity

To assess the effect of warming on leaf and root fungal colonization and alpha diversity, we used general linear mixed models. Warming treatment was included as a fixed effect and plant species and plot, both nested within warming experiment, as random effects. Alpha diversity was calculated using Hill numbers (Chao et al., 2014) with the HillR package (Li, 2018) where the “effective species number” is calculated at *q* = 0 (species richness), *q* = 1 (proportional to Shannon index), and *q =* 2 (proportional to inverse Simpsons index). As root colonization was measured as a percentage and is bounded by 0 and 1, warming effects on root colonization were tested using beta regression with the betareg package (Cribari-Neto and Zeileis, 2010). Warming effects on leaf and root alpha diversity and colonization were tested using the glmmTMB package (Brooks et al., 2017). A significant result for this test supports the hypothesis that warming alters plant-fungal endophyte symbiosis. Similarly, we ran separate models with plant species to evaluate differences among plant species in their fungal endophyte colonization and alpha diversity.

### Data analysis: Community and metabolome composition

To assess how warming and plant tissue influenced fungal endophyte community and metabolome composition, we performed multivariate distance-based redundancy analysis (dbRDA) on quantitative Jaccard distance matrices of hellinger-transformed fungal OTU tables and unit-transformed metabolome profiles using the VEGAN package (Oksanen et al., 2013). Models were conditioned using plant species and plot, both nested with warming experiment location. To test for differences among plant species or plant tissues in the magnitude of warming effects on community or metabolome composition, we performed separate dbRDA tests for each warming location and plant tissue, with plant species x warming treatment as fixed effects and models conditioned on plot. A plant species x warming interaction supports the hypothesis that warming effects significantly differed among plant species. Beta diversity of fungal endophyte communities and metabolome profiles were visualized using subsequent ordination from these models with the first two canonical dbRDA axes. To determine warming effects on the relative abundances of fungal genera, we used DESeq2 (Version 3.18; Love et al., 2014) as this approach accounts for multiple comparisons and overdispersion among taxonomic count numbers. DESeq2 was used to quantify the log2 fold response of fungal taxa to warming, aggregated based on count numbers of fungal OTUs at the genus or next lowest level of taxonomic identification.

### Data analysis: Warming effects across diverse conditions

To test whether warming effects on colonization, alpha diversity, and beta diversity differed across warming experiments and plant tissues, we used the relative interaction index (RII; Armas et al., 2004). We used RII, as opposed to other effect size estimated like log-response ration, because a) it is bounded by 1 and -1; b) is symmetrical around zero; and, c) can be calculated with variable measurements equaling zero, as is common with fungal colonization measurements (Hedges et al., 1999, Kivlin et al., 2013). We calculated the RII for these variables using the equation:

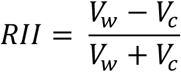

Where V_w_ is the value of the variable (colonization or diversity) in the warmed plot and V_c_ is the value of that variable in the control plot of the same block. RII was calculated for the roots and leaves of each plant species in each block. For beta diversity (community turnover), we used the Jaccard distance between the warmed and control fungal endophyte communities of plant leaves and roots for each species in each block as V_w_ and used the average Jaccard distance among leaf or root fungal endophyte communities in all control plots for each plant species as V_c_. We used generalized linear models to test whether warming effects (RII) on fungal endophytes colonization, alpha diversity, and beta diversity differed between plant tissues and among warming experiments. Plant tissue, warming experiment, and their interaction were included as fixed effects in these models. A significant plant tissue x warming experiment interaction supports the hypothesis that warming effects on fungal symbionts differ across biotic and abiotic conditions.

### Data analysis: Fungal endophyte influence on host metabolism

To test how warming influences the relationship between fungal endophyte community composition and metabolomic composition, we used distance matrix regression within each experiment. We assessed the pairwise quantitative Jaccard distance of metabolome profiles with fungal endophyte community distance, warming treatment, plant tissue, and all interactions as fixed effects. A significant fungal endophyte x warming interaction supports the hypothesis that the relationship between fungal endophytes and host metabolism is disrupted by warming.

## Results

(1) Does warming disrupt relationships between plants and fungal endophytes across roots and leaves?

### Colonization

Colonization of plant tissue by septate fungi (e.g., DSE, decomposers, and pathogens) decreased under warming in both leaves and roots (Figure 2 A, B). Septate hyphal colonization of leaves decreased 90% under warming (χ*^2^ = 10.2, p = 0.001*) and root septate colonization decreased 35% under warming (χ^2^ *= 5.33, p = 0.02*). In contrast, warming treatments did not significantly alter root colonization by aseptate (e.g., AM fungal) hyphae, arbuscules, or vesicles (Figure S1). Plant species differed in mean colonization of roots, but not leaves, with *Festuca thurberi* (RMBL) having the highest root colonization and *Elymus canadensis* (Spindletop) having the lowest (Table S1).

**Figure 2.**
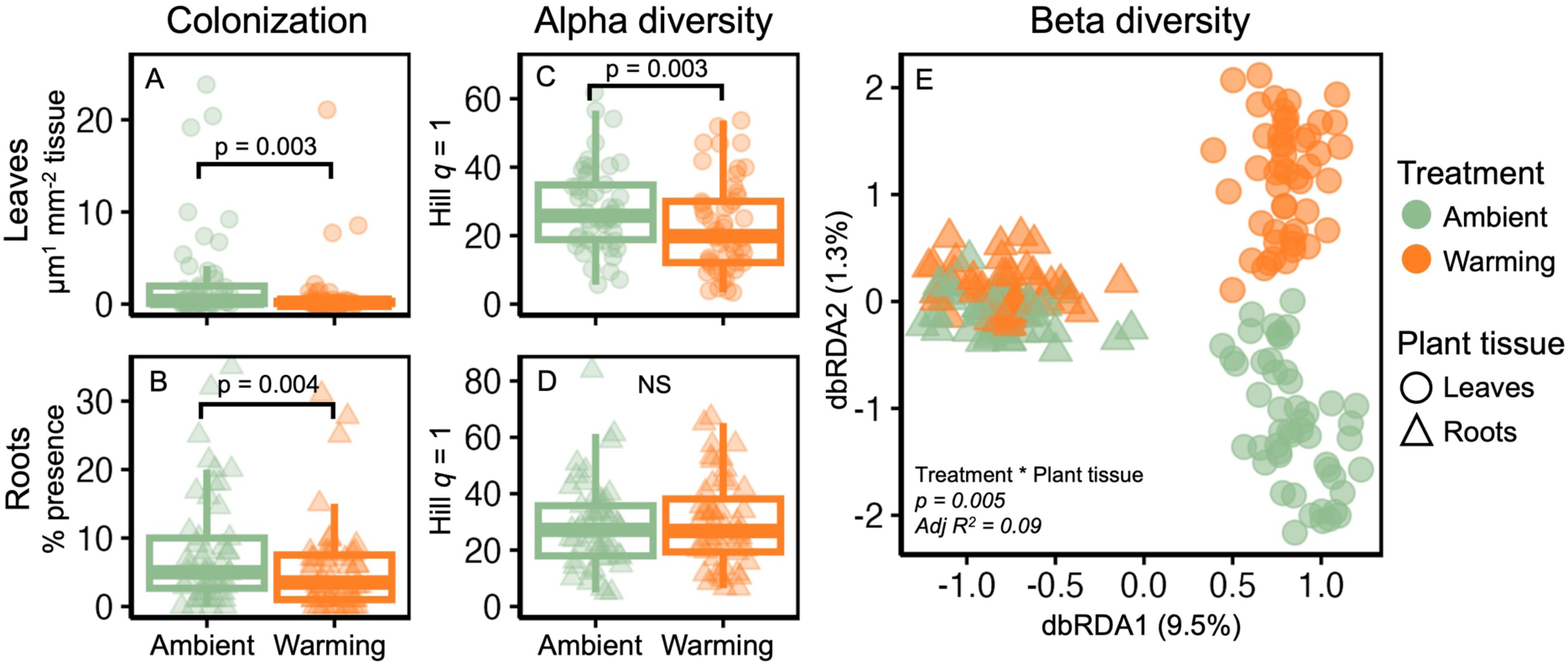
Fungal endophyte colonization (A, B), alpha diversity (C, D), and beta diversity (E) of leaf (A, C) and root (B, D) plant tissues for ambient and warming treatments across all three warming experiments. Leaf colonization rate was measured as μm septate hyphae per mm^2^ of leaf tissue while root colonization rate was measured was % root colonization by septate hyphae. Alpha diversity of leaf and root fungal endophytes was measured based on the effective number of species (Hill #) at *q* = 1 (proportional to Shannon index). Beta diversity was assessed as Jaccard dissimilarity among fungal endophyte communities. Beta diversity ordination was constructed based on distance-based redundancy analysis (dbRDA) with treatment, plant tissue, and their interaction as fixed effects, constrained by plant species and plot, both nested within warming experiment location. The variation explained by the first two dbRDA axes from these models is shown in parenthesis on the axis.

### Diversity

Fungal endophytic alpha diversity decreased under warming in plant leaves, but not roots (Figure 2 C, D). In plant leaves the effective species number was 20% lower (*χ^2^ = 8.67, p = 0.003*) at Hill *q* = 1 (Shannon index), 11% lower (*χ^2^ = 4.03, p = 0.04*) at Hill *q* = 0 (species richness), and 21% lower (*χ^2^ = 7.70, p = 0.006*) at Hill *q* = 2 (inverse Simpson index) in warmed compared to ambient leaves (Figure S2). Although roots had higher diversity than leaves across all orders of *q* (Figure 2 C, D), warming did not significantly reduce the alpha diversity of endophytes in roots. Plant species also varied in leaf and root alpha diversity (Table S1). *Sorghastrum nutans* (Kessler) and *Poa pratensis* (RMBL) had the highest alpha diversity (*q* = 1) in their leaves and roots, respectively. *Elymus canadensis* and *Sporobolus compositus* had the lowest alpha diversity (*q* = 1) in their leaves and roots, respectively.

### Fungal endophyte composition

Warming shifted fungal endophyte communities, but these effects were stronger in leaves than in roots. Across all samples, the effect of warming varied between leaves and roots (*p = 0.005*, Figure 2E), with this interaction explaining 9% of variance in fungal endophyte community composition. Warming treatment explained 1.8% of variance in leaf community composition (*p = 0.001*) versus 0.1% of root community composition (*p = 0.03*).

One hundred two fungal genera significantly responded to warming (Figure 3), with 76% of responding genera occurred in leaves versus 24% in roots. Of responding genera, 42% were identified as saprotrophs, 27% were pathogens, and 6% were mutualists. The majority of responding genera (67%) declined with warming, particularly in the mutualists and pathogens. There were 8 responding genera that occurred in both leaves and root, however not all responded in the same direction. The genera *Xenopenidiella* and *Pyrenochaetopsis* were both negatively affected by warming while *Marasmius*, *Mollisia,* and *Filobasidium* responded positively to warming across plant tissues. The genera *Emmonsiellopsis*, *Colletotrichum*, and *Stagonospora* were negatively affected by warming in plant leaves but responded positively to warming in roots. Most responding taxa were relatively common across samples, in over half (57%) of samples on average (min 15% and max 100%) and making up an average of 0.3% of the overall fungal community (min 0.005% and max 5.1%). Some responding taxa (25%) could not be assigned to a functional group due to missing annotations or lack of genus-level identification (Table S2). Over half of all responding genera belonged to Dothideomycetes (39%) or Sordariomycetes (14%), with Dothideomycetes being the most abundant responding class for mutualists, pathogens, and endophytes (Figure S3).

**Figure 3.**
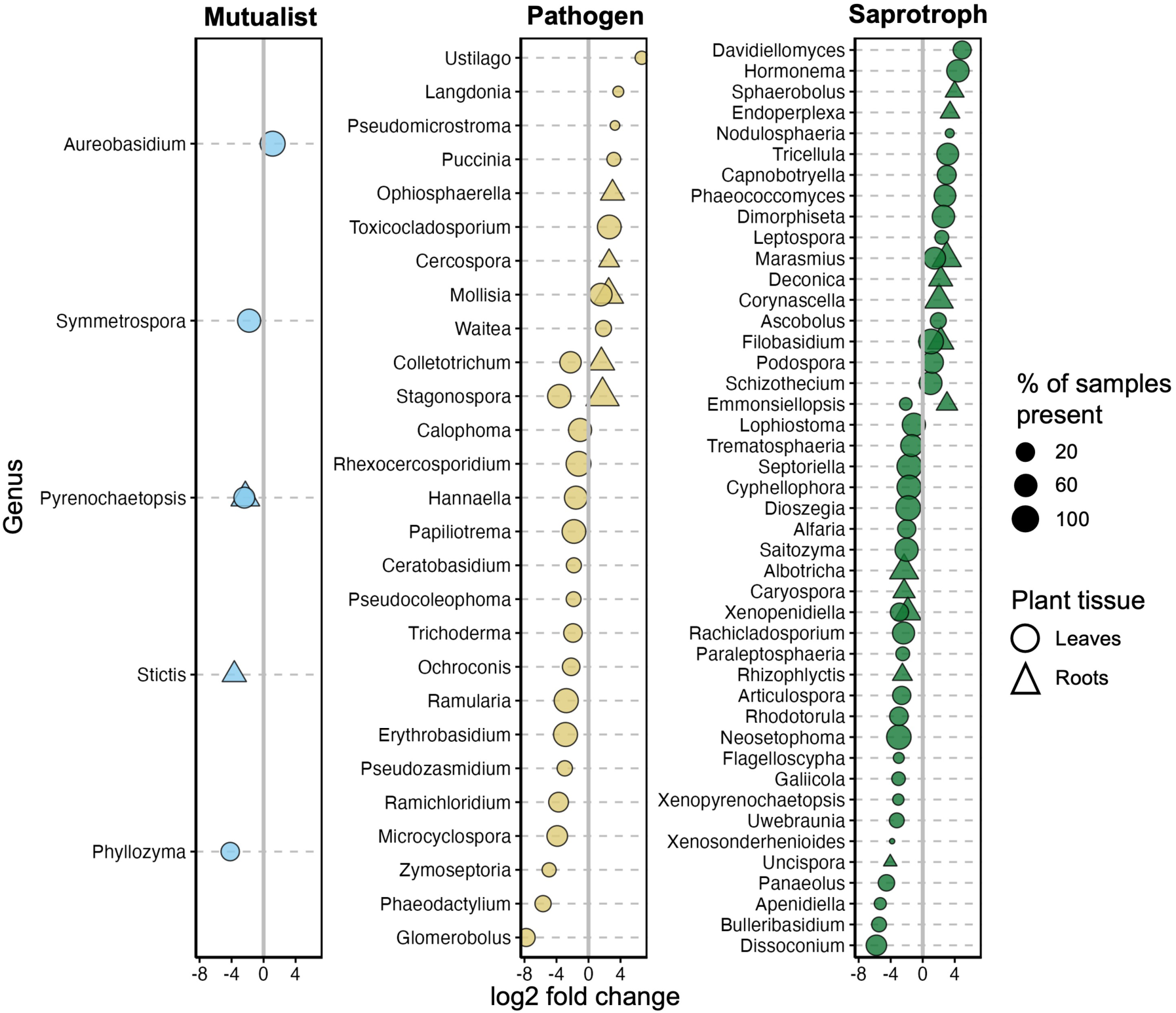
Responses of fungal symbiont genera to warming based on deseq2 algorithm where positive values represent greater abundance in warmed treatments and negative values represent greater abundance in ambient treatments. All genera shown demonstrated a significant response at adjusted p-value < 0.05. Genera were assigned to putative functional groups based on the fungal traits database. Circles represent genera with significant responses to warming (*P*<0.05) in leaves and triangles represent those with significant responses in roots.

(2) Does warming alter leaf and root fungal endophytes in similar ways across diverse environmental conditions?

The influence of plant tissue and warming experiment on fungal endophyte responses to warming differed across metrics of fungal endophyte abundance and diversity. Fungal endophyte colonization decreased with warming in both leaf and root tissues, but effects were greater in the Kessler experiment than at RMBL or Spindletop (*χ^2^ = 6.66, p = 0.02*, Figure 4 A). Across all experiments, warming decreased fungal alpha diversity in leaves, but not roots (*χ^2^ = 8.64, p = 0.003*, Figure 4 B). Warming effects on fungal endophyte beta diversity (community turnover) depended on both plant tissue and warming experiment (*χ^2^ = 20.29, p < 0.001*, Figure 4 C). Warming increased fungal endophyte community turnover in plant leaves at the Kessler and Spindletop experiments, as well as plant roots in Spindletop. However, warming did not increase turnover in plant roots at the Kessler experiment, as well as leaves and roots at the RMBL experiment (Figure S4).

**Figure 4.**
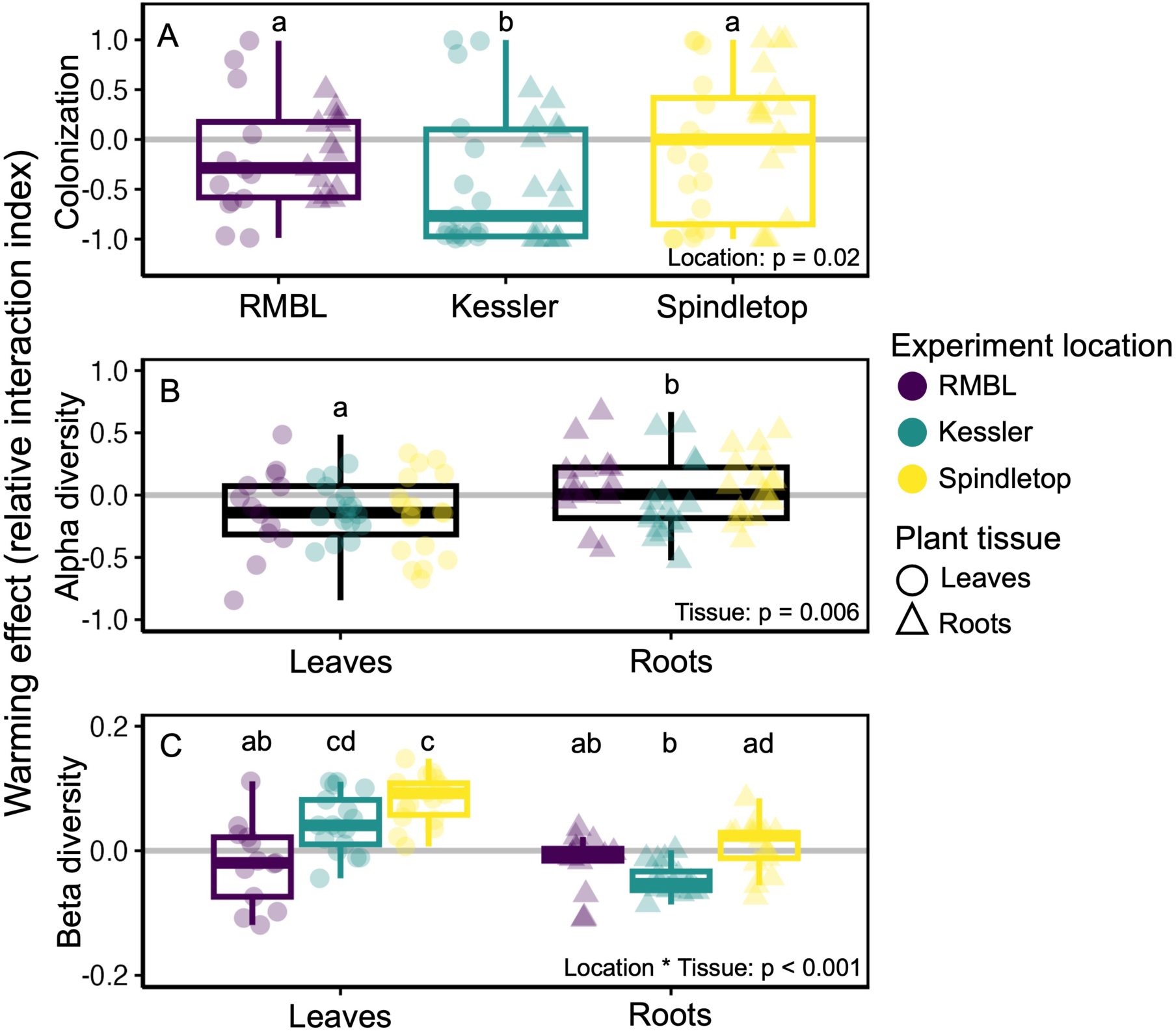
Effects of warming (relative interaction index) on fungal endophyte colonization (A), alpha diversity (B), and beta diversity (community turnover, C) in plant leaves and roots across the Rocky Mountain Biological Station (RMBL), Kessler, and Spindletop warming experiments. A positive warming effect indicates warming increased this variable, while a negative effect indicated warming decreased this variable. Groups with the same letter within each variable are not significantly different from each other at p < 0.05.

(3) Do warming-induced changes to fungal endophyte communities correspond to altered host function via plant metabolic activity?

Warming weakened the relationship between fungal endophyte composition and metabolomic composition in both leaf and root tissues (endophyte x warming: *χ^2^ = 7.*71, p = 0.005, Figure 5). Warming did not strongly affect plant metabolome profiles, though metabolomes differed significantly between plant tissues and among plant species (*p < 0.05*, Figure S5). In plant leaves, warming decreased the correlation between fungal endophyte and metabolome composition by 70% (ambient β = 0.41; warming β = 0.12; Figure 5 A, B). In plant roots the relationship between endophytes and metabolomes was weaker but still demonstrated the same overall response to warming as in leaves with a decrease of 80% (ambient β = 0.15; warming β = 0.03; Figure 5 C, D).

**Figure 5.**
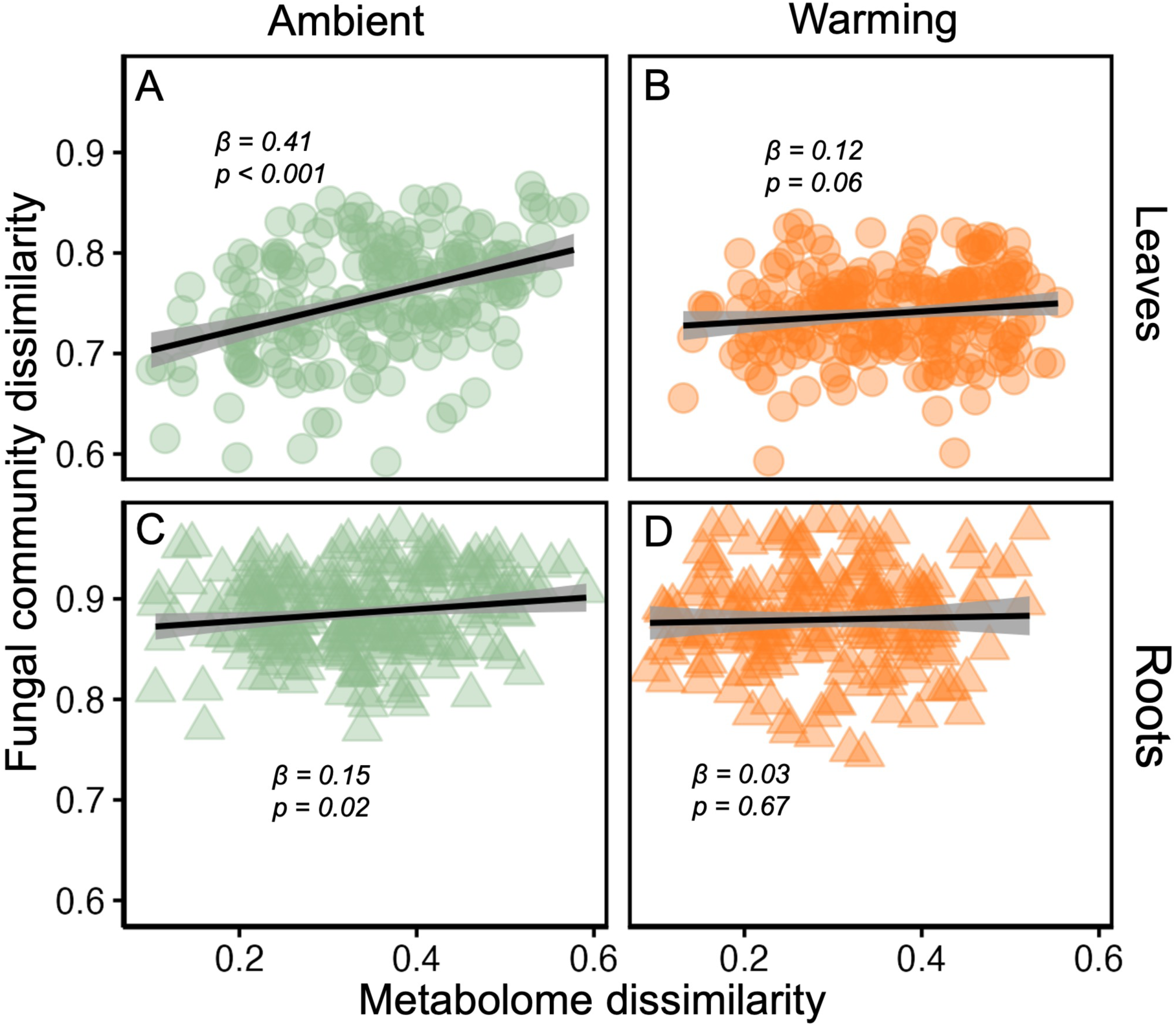
Correlations between fungal endophyte community and metabolome dissimilarity for leaf (A, B) and root (C, D) plant compartments in ambient (A, C) and warming (B, D) treatments. Values represent pairwise Jaccard distances among all samples within the RMBL and Kessler experiments.

## Discussion

Warming generally decreased the strength of interactions between plants and fungal endophytes. Warming effects on fungal endophytes were more pronounced in leaves than roots and differed in magnitude among ecosystems that had different plant species, climates, and experimental warming treatments. Importantly, warming disrupted the relationship between fungal endophyte composition and plant metabolomic composition, suggesting fungal endophytes may contribute less to plant physiology under climate warming. As plants adapt to cope with the stresses of increasing temperature, they may alter resource use patterns (Crous, 2019) through shifts away from fungal endophytes and in turn receive less direct benefit. Overall, warming-driven disruption of fungal endophyte community composition and function shows the sensitivity of this symbiosis to global change. This poses concerns for the role of fungal endophytes in plant stress amelioration and promoting plant resilience to the degree of warming projected under most climate change models (Canadell et al., 2023).

### Does warming disrupt relationships between plants and fungal endophytes by reducing colonization and diversity or altering community composition of fungal endophytes, and are effects greater for leaves than roots?

Warming caused colonization of plant tissues by septate fungi to decrease across leaves and roots. However, warming-induced reductions to fungal endophyte diversity and changes to community composition were restricted to leaves. Warming temperatures tend to shift environmental conditions outside of optimal ranges for native fungal endophytes, negatively affecting these communities (Bálint et al., 2015, Randriamanana et al., 2015). These changes are often reflected in declines in the diversity and evenness of fungal endophytes (Faticov et al., 2021) as well as overall compositional changes in communities (Lyons et al., 2021) that can favor potentially harmful fungal taxa (Sui et al., 2020). However, warming can also increase fungal colonization of plant tissues as hosts rely more on fungal symbionts to overcome increased thermal stresses (Olsrud et al., 2010, Rudgers et al., 2014, Staddon et al., 2004). Notably, septate colonization of root tissues of *Lolium arundinaceum* was previously found to increase with warming at one of the warming experiments used here (Spindletop; Slaughter et al., 2018), though a different cultivar. While some plant species or cultivars may respond positively to warming, our results support the finding of a previous metanalysis (Kivlin et al., 2013) that warming has a generally negative effect on fungal endophytes. This is particularly true for leaves, where warming also decreased fungal endophyte diversity and altered community composition, but may be more complicated for roots. We did not find decreases to fungal diversity in roots from warming, nor did we find any changes to the colonization of arbuscular mycorrhizal fungi. Warming can stimulate carbon allocation to arbuscular mycorrhizal fungi (Rillig et al., 2002) but may also reduce available root area to colonize (Qiu et al., 2021). Soil properties can mitigate the impacts of global change factors on belowground communities (Fridley et al., 2011), such as buffering soil dwelling fossorial animals from increased temperatures (Cameron and Scheel, 2001). Further, changes to root symbionts following warming can be better attributed to altered host performance than direct products of climate change (Fernandez et al., 2017). These patterns demonstrate that warming will likely cause a quantitative decline in the strength of interactions between plants and endophytic fungi, with the effects more apparent in leaves than roots as hosts drive these changes indirectly potentially due to soil buffering (Jiang et al., 2021).

While fungal endophytes can perform many different functional roles (*i.e.,* decomposers, pathogens, mutualists), most beneficial fungal endophytes in plant tissues are septate, indicating potential declines in benefits to plants with warming. Across all functional groups, more genera of fungal endophytes responded negatively to warming than positively, but this was more acute for putatively symbiotic taxa (mutualists and pathogens) than commensal (saprotrophs). We found that Dothideomycetes and Sordariomycetes (both Ascomycota) were the most abundant classes among fungi responding to warming, with both negative and positive responses depending on genus. Taxon-specific responses to warming often show that Ascomycota are relatively sensitive to warming (Xiong et al., 2014, Jiang et al., 2021). However, based on the inconsistency of their responses, taxonomic relationships may be most reliable to predict which fungi might be impacted by warming, but not the direction of their response. Interestingly, the most abundant leaf mutualist was *Aureobassidium*, a yeast-like fungi that can help confer disease resistance (Pinto et al., 2018). This genus increased in abundance due to warming. Though warming will likely have an overall negative effect on symbiosis between plants and fungal endophytes, this finding suggests that some positive interactions could persist to promote plant resilience. Conversely, while pathogens were also largely disrupted by warming (Chen et al., 2024), some common genera of pathogens (*e.g., Ustilago)* were greatly increased on leaves with warming. Prevalent pathogens that can withstand warmer temperature, like leaf smuts (Raza and Bebber, 2022), may pose greater risks to plants under future climates. Overall, the sensitivity of fungal endophyte communities, particularly in leaves, to global change could highlight further areas of future vulnerability for plant health and ecosystem function (Berg and Cernava, 2022). Yet, if we can identify mutualists with positive responses to warming, these endophytes could provide potential avenues to increase plant resilience (Suryanarayanan and Shaanker, 2021).

### Does warming alter leaf and root fungal endophytes in similar ways across diverse environmental conditions?

Warming generally altered fungal endophyte communities across all experimental conditions. However, these changes were not uniform. Environmental filtering features greatly in fungal endophyte community assembly (Pellitier et al., 2019, Ricks and Koide, 2019) by geography (Harrison and Griffin, 2020), climate (Giauque and Hawkes, 2013), edaphic conditions (Glynou et al., 2016), and plant hosts (Christian et al., 2016). Given these sources of potential variation in endophytic response to warming, it is important to delineate consistent processes from context-specific patterns of community assembly. The design of our study confounded many of the factors due to the idiosyncratic nature of individual warming experiments. Thus, multiple, co-varying differences among experiments could underlie observed variation in endophytic response to warming. These differences include warming intensity, length of time since warming, as well as biotic and abiotic factors (*e.g.,* surrounding communities, soil types, distance to other anthropogenic disturbances, etc.). Our study showed that both biotic and abiotic factors are important determinants of endophytic response to warming, but more evidence is needed to better estimate the independent and synergistic effects of these components.

The greatest changes to endophyte community composition were observed in the most recent warming experiment (Spindletop), while the oldest warming experiment (RMBL) generally had the least changes to endophyte communities. These temporal patterns could be indicative of thermal acclimation in local fungal communities (Crowther and Bradford, 2013). The experiments occurred over a temporal range of 25 years, but the temperature range was only 2 °C. Temporal fluctuations (Faticov et al., 2021), tipping points (Jassey et al., 2018), or plant/fungal acclimation (Perreault and Laforest-Lapointe, 2022) in endophytic response to warming could lead to alternative patterns observed over time as ecosystems warm at different rates. In our experiment, the endophytes in grass species under >20 years of warming shifted less in diversity and composition compared to the more recent (<10 years) warming experiments, suggesting that acclimation might have required decadal time periods.

The Kessler experiment, which comprised grasses with C_4_ photosynthetic pathways, experienced the greatest overall reduction in fungal endophyte colonization. As in many other studies, host-species identity was one of the strongest predictors of endophytic community composition (Kivlin et al., 2022) and thus also likely represents a large source of plasticity in plant-fungal response to global change (Wrzosek et al., 2017). Warming is expected to promote the growth of C_4_ grasses due to their ability to conserve water (Morgan et al., 2011), however it can also greatly alter their resource allocation, foliar chemistry, and stoichiometry across plant tissues (Habermann et al., 2019). These changes can negatively impact the fungi residing in their tissues (de Oliveira et al., 2020), despite their potential benefit for the plants. As C_4_ grasses dominate in regions expected to be particularly vulnerable to global change (Edwards and Still, 2008), their increased sensitivity to symbiotic disruption could exacerbate these effects.

### Does warming decouple fungal endophyte communities from plant metabolic activity?

We predicted that plant response to warming would disrupt the relationship between plant metabolism and fungal endophyte composition. Warming may disconnect fungal endophytes from plant metabolic regulation based on the result that coordination between fungal endophyte communities and metabolomic profiles weakened under warming for both leaves and roots. Fungal endophytes can strongly influence host plant phenotype via production of secondary metabolites (Sandy et al., 2023). Warming-induced shifts in plant metabolomic profiles can occur without genetic changes in hosts (Sun et al., 2022), suggesting alternative drivers such as symbiotic microbes. Despite disruption between fungal endophytes and host metabolism, we found no overall differences in metabolomic profiles in response to warming. Limited independent impacts of increased temperatures on plant metabolomes have been reported previously (Gargallo-Garriga et al., 2015). This suggests that plants may maintain metabolic homeostasis under warming (Dusenge et al., 2019). However, such maintenance may occur at the expense of other functions, such as pathogen suppression (Liu and He, 2021), drought tolerance (Gargallo-Garriga et al., 2015), or reproductive output (Liu et al., 2012). These synergistic contributors toward decoupling of plants and their symbiotic microbes under warming are likely to exacerbate the impacts of other global change stressors (Zhu et al., 2022). The functional role of fungal endophytes in host fitness will be critical to forecast plant responses to warmer conditions (Naik, 2019).

## Conclusions

Fungal endophytes have been proposed as a potential biological resource to help plants acclimate to climate change. Our results show that the strength of plant-fungal symbioses will likely decline with climate warming. Although negative effects of warming mostly occurred for leaf endophytes, effects were not consistent across experimental conditions or plant species. These context-dependent responses to warming suggest that conservation efforts should target locations where endophytes are most vulnerable and could identify genetic resources to increase plant and fungal resilience. However, our results suggests that fungal endophytes may not be a significant mechanism for enhanced plant resilience to global change.

## Supporting information

endophyte_supplemental

## Acknowledgements

JDE was supported by the U.S. National Science Foundation Division of Environmental Biology grant #2305863. SNK was supported by NSF grants DEB2217353, DEB 2106065, and DEB1936195 and the U.S. Department of Energy, Office of Science, Office of Biological and Environmental Research, Terrestrial Ecosystem Sciences program under award number DE-FOA-0002392. JAR was supported by NSF#1354972. RLM was supported by U.S. Department of Energy (08-SC-NICCR-1073), NSF (DEB1021222), the Kentucky Agricultural Experiment Station (KY006045), and a cooperative agreement with the USDA-ARS Forage Animal Production Research Unit (58-6440-7-135).

