## Supplementary material for "Warming disrupts plant-fungal endophyte symbiosis more strongly in leaves than roots": endophyte_supplemental

**Table S1**. Species mean for fungal endophyte colonization and alpha diversity in leaves and roots with standard error (in parentheses). Post-hoc differences for variables with species differences are shown with letters such that species with the same letter are not significantly different from each other at p < 0.05 for that variable.

| Warming experiment | RMBL | | | Kessler | | | Spindletop | | | |
|---|---|---|---|---|---|---|---|---|---|---|
| Species | *Achnatherum lettermanii* | *Festuca thurberi* | *Poa pratensis* | *Schizachyrium scoparium* | *Sorghastrum nutans* | *Sporobolus compositus* | *Elymus canadensis* E- | *Elymus canadensis* E+ | *Lolium arundinaceum* E- | *Lolium arundinaceum* E+ |
| Leaf colonization | 0.37 (0.04) | 0.99 (0.27) | 1.02 (0.18) | 4.31 (0.61) | 5.8 (0.95) | 1.66 (0.28) | 0.32 (0.04) | 0.38 (0.05) | 0.5 (0.08) | 1.2 (0.25) |
| % septate root colonization | 7.7 (0.58) ab | 15.11 (1.16) a | 5.87 (0.74) ab | 11.08 (1.12) ab | 4.75 (0.61) b | 4.56 (0.61) b | 3.09 (0.32) b | 5.87 (0.32) ab | 3.86 (0.39) b | 5.04 (0.32) b |
| % aseptate root colonization | 66.46 (1.47) a | 59.59 (1.86) ab | 45.35 (1) abc | 36.3 (2.07) bc | 28 (1.49) c | 29.33 (2.33) c | 48.97 (1.27) abc | 47.1 (1.59) abc | 39.01 (1.26) bc | 38.75 (2.54) bc |
| % arbuscule root colonization | 4.29 (0.39) | 4.26 (0.63) | 1.99 (0.23) | 3.07 (0.38) | 1.74 (0.33) | 1.69 (0.27) | 3.32 (0.32) | 2.66 (0.22) | 4.21 (0.42) | 2.55 (0.21) |
| % vesicle root colonization | 6.19 (0.37) ab | 8.01 (0.56) a | 4.54 (0.36) abc | 2.57 (0.25) bc | 1.25 (0.17) c | 1.79 (0.19) bc | 3.97 (0.31) abc | 4.65 (0.33) abc | 5.66 (0.25) abc | 4.87 (0.38) abc |
| Leaf alpha q0 | 380.44 (6.8) ab | 543.3 (22.06) abc | 558.75 (22.88) abc | 615.5 (14.14) c | 472.58 (13.77) abc | 556.18 (16.12) ac | 444 (13.87) abc | 647.44 (14.45) ]c | 340.3 (13.19) b | 380.1 (14.82) ab |
| Leaf alpha q1 | 14.46 (0.62) ab | 20.14 (0.96) abc | 22.04 (1.44) abc | 32.37 (1.1) cd | 40.84 (1.33) d | 31.71 (1.19) cd | 11.31 (0.73) a | 29.36 (0.68) bcd | 21.29 (0.6) abc | 20.66 (0.93) abc |
| Leaf alpha q2 | 4.98 (0.23) a | 6.29 (0.32) ab | 5.92 (0.36) ab | 12.88 (0.47) cd | 17.84 (0.78) c | 11.95 (0.51) bd | 4.58 (0.3) a | 12.58 (0.4) bcd | 8.66 (0.29) abd | 8.52 (0.41) abd |
| Root alpha q0 | 659.2 (25.04) a | 633.1 (20.67) ab | 574.78 (27.38) ab | 557.42 (19.34) ab | 415.58 (12.13) b | 415.5 (13.69) b | 612.11 (19.88) ab | 684.78 (23.21) a | 737.44 (23.96) a | 796.1 (25.77) a |
| Root alpha q1 | 32.64 (1.08) abc | 35.38 (2.07) ab | 40.55 (1.2) a | 24.18 (1.28) bcd | 17.56 (0.75) cd | 15.6 (0.84) d | 25.87 (0.75) bcd | 22.79 (0.85) bcd | 39.53 (1.62) ab | 37.85 (1.34) ab |
| Root alpha q2 | 12.9 (0.49) abc | 11.87 (0.8) abc | 14.85 (0.59) ab | 9.42 (0.48) ac | 7.28 (0.3) c | 6.36 (0.3) c | 9.06 (0.31) abc | 6.49 (0.24) c | 16.48 (0.84) b | 14.41 (0.49) ab |

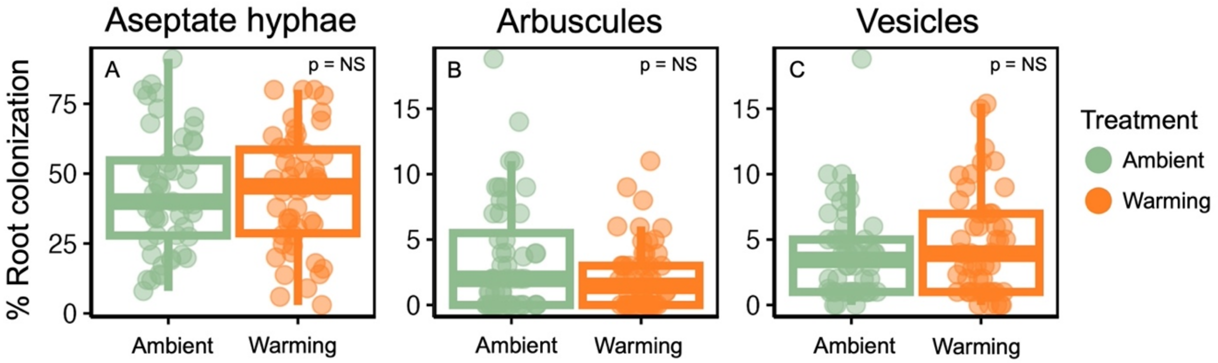


Figure S1. Root colonization rates for ambient and warming treatments representing % root colonization of aseptate hyphae (A), arbuscules (B), and vesicles (C) associated with arbuscular mycorrhizal fungi.

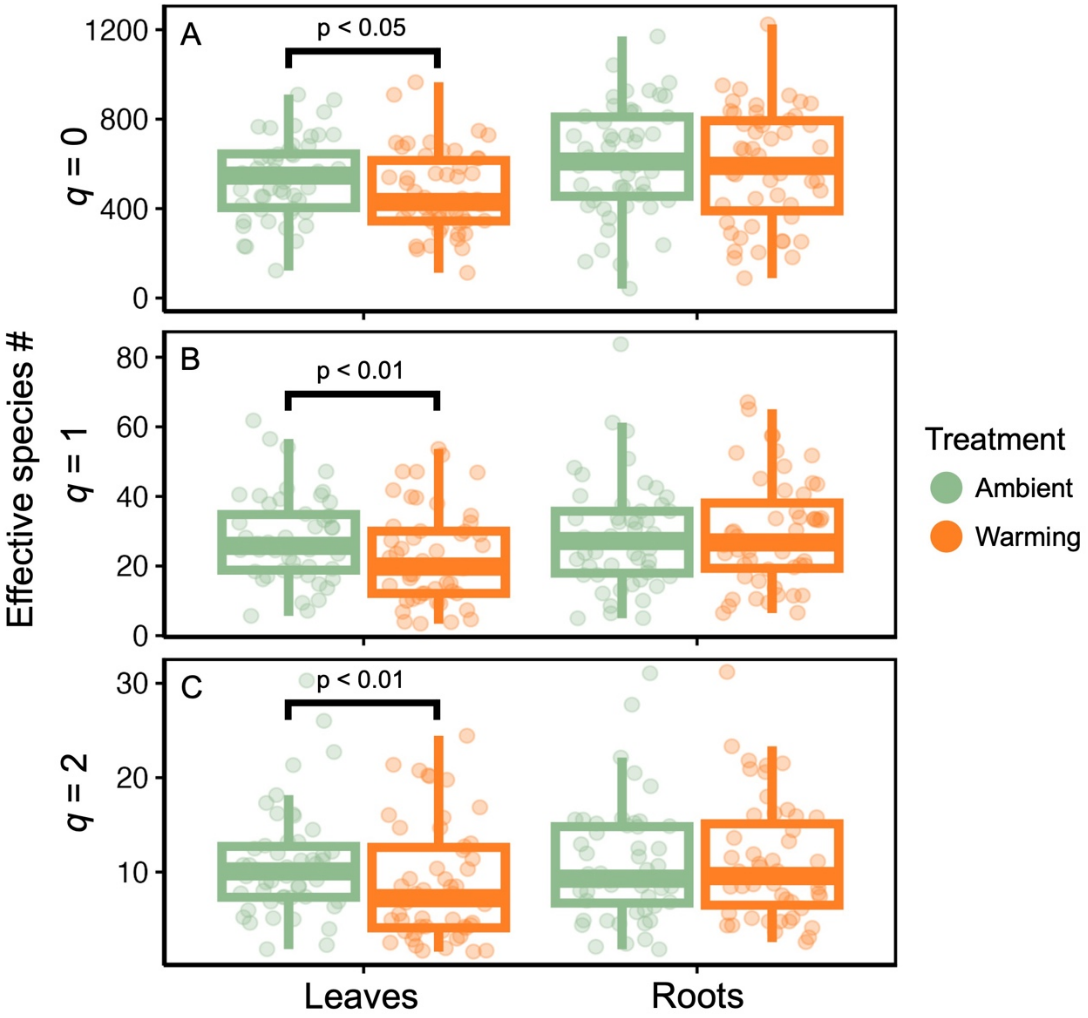


Figure S2. Alpha diversity of fungal endophytes in leaf and root compartments of warming treatments with Hill numbers depicting effective number of species with all OTUs weighted equally ($q$ = 0; A), OTUs weighted based on their proportional abundance ($q$ = 1; B), and rare OTUs down-weighted ($q$ = 2; C).

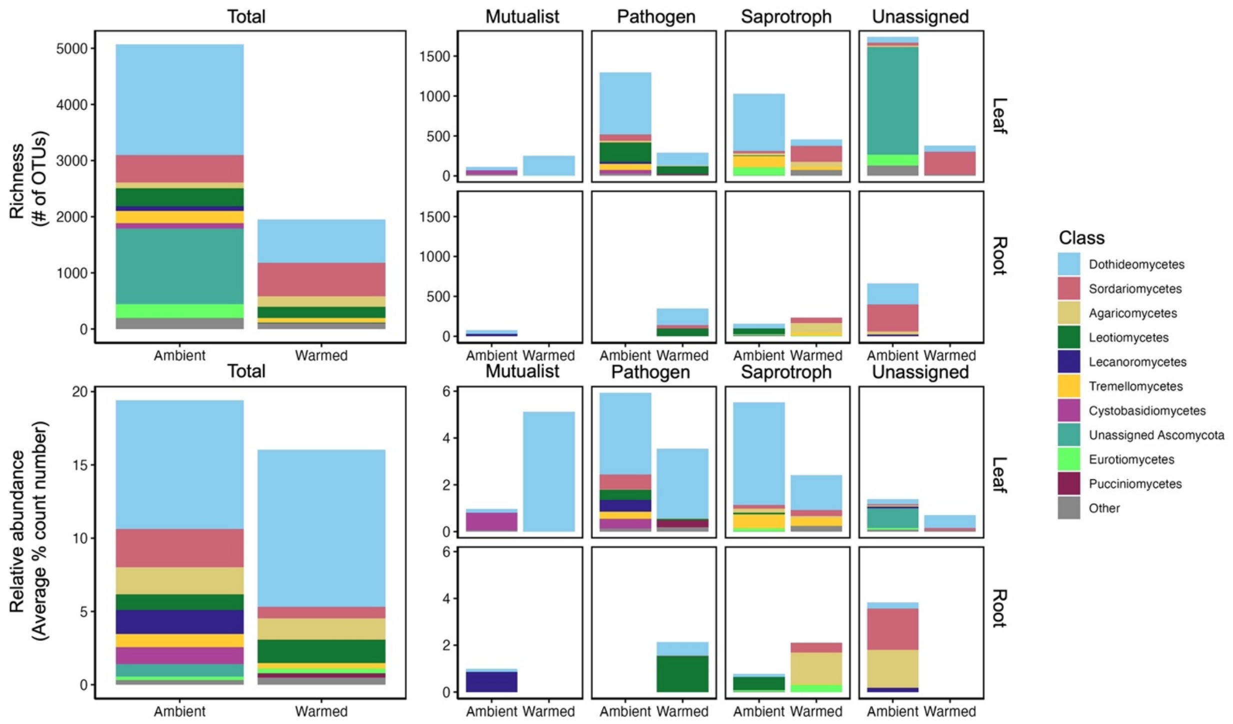


Figure S3. Richness (# of OTUs) and relative abundance (average % count number) of genera with a significant (adjusted $p < 0.05$) response to warming based on DeSeq2 analysis, depicted with class-level assignments, shown for both the total fungal endophyte community and the functional group x plant tissue specific combinations.

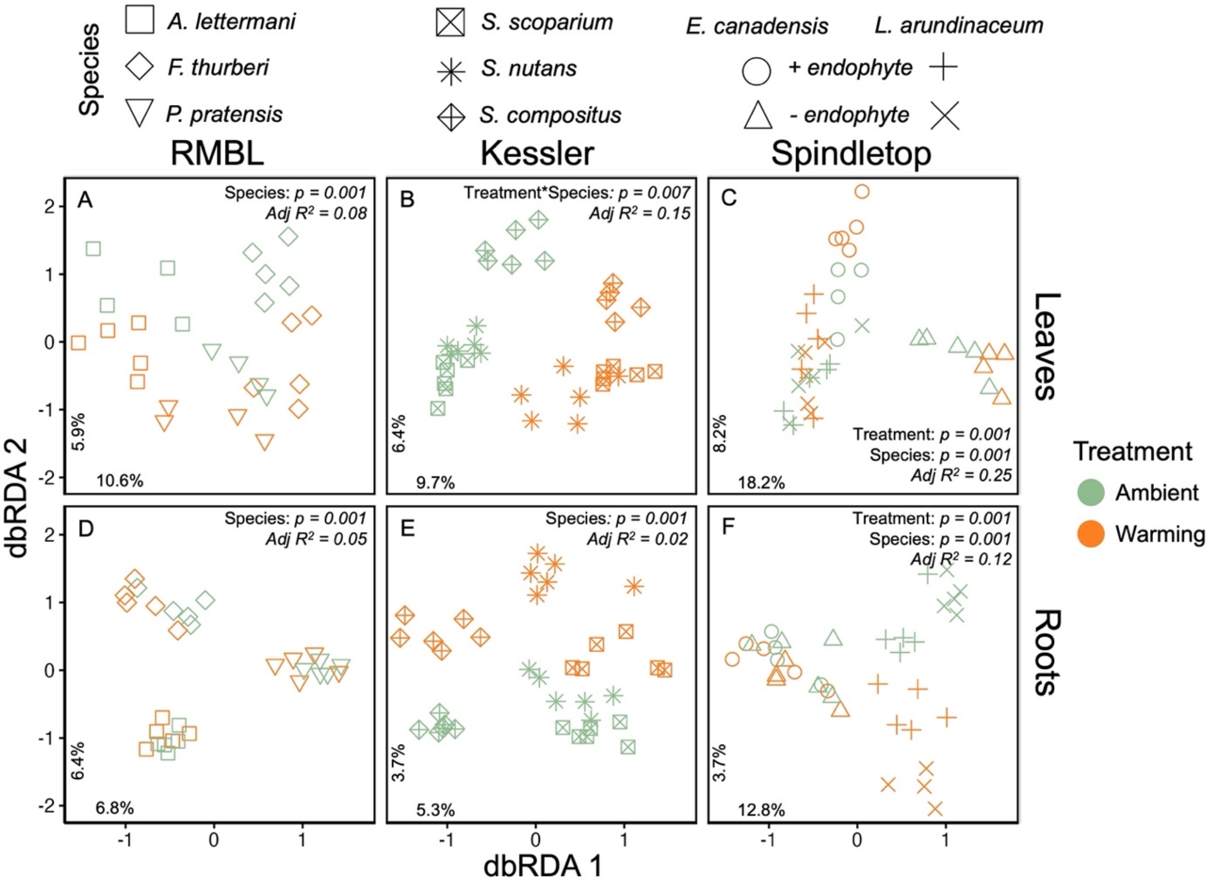


Figure S4. Fungal endophyte community ordinations for leaves (A-C) and roots (D-E) in ambient and warming treatments of each species at each warming experiment; Rocky Mountain Biological Laboratory (RMBL; A, D), Kessler (B, E), and Spindletop (C, F) . Ordinations were constructed based on distance-based redundancy analysis (dbRDA) models using treatment, species, and their interaction as fixed effects, constrained by experiential plot. The variation explained by the first two dbRDA axes from these models is shown in the lower left-hand corner of each ordination.

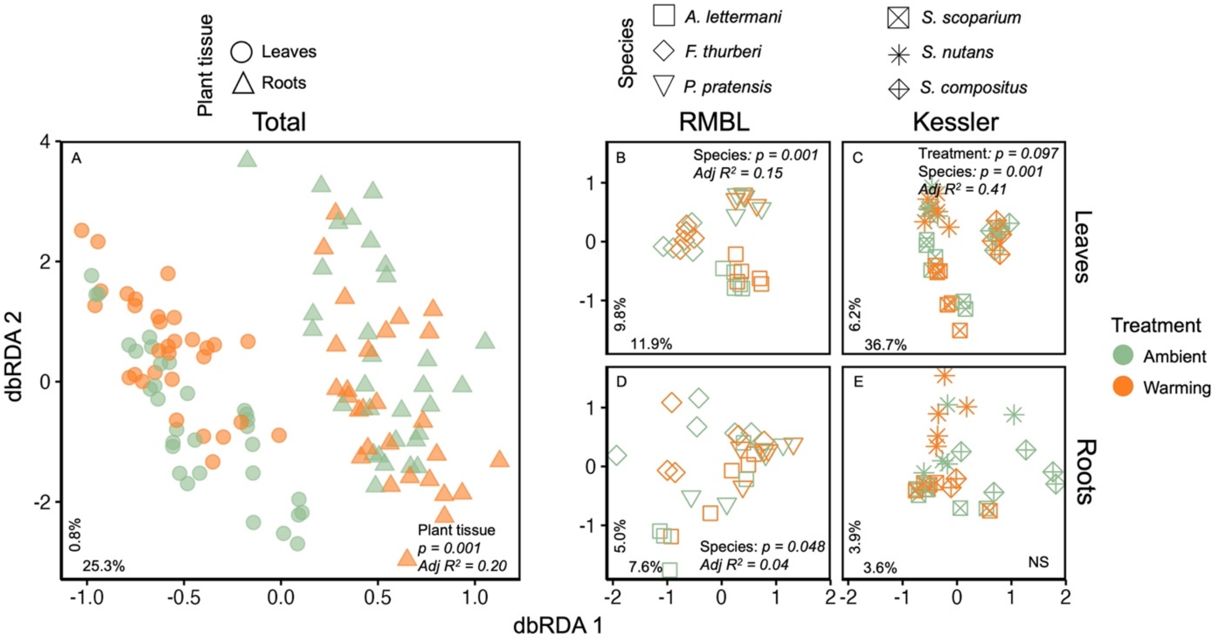


Figure S5. Metabolome dissimilarity profile odinations in ambient and warming treatments for the total metabolome (A) and for leaves (B, C) and roots (D, E) of each species at the Rocky Mountain Biological Laboratory (RMBL; B, D) and Kessler (C, E) warming experiments. Ordinations were constructed based on distance-based redundancy analysis (dbRDA) models using warming treatment, plant tissue, and their interaction as fixed effects constrained by species and experimental plot nested within warming experiment site for total metabolome (A) and warning treatment, plant species, and their interaction as fixed effects, constrained by experiential plot for warming experiment x plant tissue specific plots (B - E). The variation explained by the first two dbRDA axes from these models is shown in the lower left-hand corner of each ordination.
